# Enumerating all possible biosynthetic pathways from metabolic networks

**DOI:** 10.1101/226795

**Authors:** Aarthi Ravikrishnan, Meghana Nasre, Karthik Raman

## Abstract

Exhaustive identification of all alternate possible pathways that exist within metabolic networks can provide valuable insights into cellular metabolism. With the growing number of metabolic reconstructions, there is a need for an efficient method to enumerate pathways, which can also scale well to large metabolic networks, such as those corresponding to microbial communities.

We developed MetQuest, an efficient graph-theoretic algorithm to enumerate all possible pathways of a particular length between a given set of source and target molecules. Our algorithm employs a *guided* breadth-first search to identify all feasible reactions based on the availability of the precursor molecules, followed by a novel dynamic-programming based enumeration, which assembles these reactions into pathways producing the target from the source. We demonstrate several interesting applications of our algorithm, ranging from predicting amino acid biosynthesis pathways to identifying the most diverse pathways involved in degradation of complex molecules. We also illustrate the scalability of our algorithm, by studying larger graphs such as those corresponding to microbial communities, and identify several metabolic interactions happening therein.

A Python-based implementation of MetQuest is available at https://github.com/RamanLab/MetQuest

## 1 Introduction

Genome-scale metabolic networks are very useful to understand the complex network of metabolic reactions happening inside cells [37]. Typically, a genome-scale metabolic network consists of thousands of reactions and metabolites, which capture vital metabolic pathways such as the biosynthesis of amino acids and lipids, ATP synthesis, as well as transport of molecules inside the cells. Over the recent years, the construction of such networks has increased tremendously, in part also due to the growing number of genomes sequenced [34]. There are several well-established constraint-based methods to analyse metabolic networks [41]; however, these methods require well-curated genome-scale metabolic models for making reliable predictions. The number of such well-curated models are very few [13, 34, 42], in comparison to the genome sequences available. Alternative approaches to analyse genome-scale models are based on network “topology” [4, 5, 18] or Boolean rules [26, 27]. In the former, the metabolic networks are abstracted as networks or graphs, and graph-theoretic algorithms are employed to predict or infer metabolic pathways. The latter set of methods are based on definitions of Boolean functions for identifying if particular reactions can proceed or not. Both methods generate qualitative predictions and do not entail the requirement of well-curated genome-scale models.

Several graph-based methods, using naïve network expansion [17,18], atom–atom mapping, subgraph matching or stoichiometry [4, 22, 35, 39] have been developed previously. These methods, regardless of the approach, aim to determine the route(s) of conversion between sets of source and target molecules. The algorithms based on network expansion traverse the graph depending on the availability of precursor compounds and determine different routes of conversions from the metabolic network. The input graph to these algorithms can be represented in different ways, such as a substrate graph [28] or a hypergraph [32], each of which captures information at different levels of complexity. Substrate graphs are one of the simplest forms of representation, where all metabolites participating in a reaction are connected to each other. Traversing these graphs using breadth-first or depth-first search to identify different routes of conversion often leads to erroneous paths such as a 2-step glycolysis, converting glucose to pyruvate (via ADP), since these could proceed using the connections between “side” metabolites. To avoid such spurious results, several tools, such as FMM (From Metabolite to Metabolite; [7]) and Metabolic Tinker [30] use substrate graphs after excluding the connections from (high degree) “currency metabolites” such as H^+^, Water and NADPH. However, the substrate graph representation fails to capture the requirement of more than one reactant for any given reaction to occur.

To overcome these problems, algorithms were developed to operate on more informative bi-partite and hypergraph representations of metabolic networks [10,11,17,18,32]. These representations consider the participation of multiple compounds in a reaction to produce product(s), which is crucial while depicting metabolic networks. One such hypergraph-based algorithm, Rahnuma [32], performs a depth-first search to identify pathways that lead to the target metabolite from the source. In a few other studies [17,18], the metabolic networks are represented as bipartite graphs to first determine the scope of the starting seed metabolites. The synthesis pathways were then assembled by backtracking from the target to the source, which becomes computationally challenging due to the presence of branched and cyclic pathways in metabolic networks.

The problem with determining branched-pathways has been addressed in few studies, which seek to use atom tracing. In one such method [19] BPAT-S (Branched Pathfinding using Atom Tracking and Seed pathways), *seed pathways* (linear) are first identified between the source and target compounds, based on the atom loss/gain. The branched pathways are then identified by finding the *linear pathways* between the metabolites contributing to this atom loss/gain. These pathways are then ranked based on the total number of reactions and the number of conserved atoms. ReTrace [40], another algorithm to find branched pathways, tries to combine linear shortest paths based on the transfer of a high fraction of atoms from source to target, thereby finding only the k-shortest paths.

Another class of methods use either information from atom-mapping, thermodynamics or structural transformation. For instance, the algorithm developed by [4] defines a set of atom-mapping rules based on the possible chemical conversions and seeks to identify paths where the atom loss is minimum. The predictions of this algorithm is restricted to the first *k*-shortest paths. PathPred [35], another tool to predict metabolic pathways, performs subgraph matching on the graph generated from KEGG RPAIR database. Similarly, RouteSearch [25], another algorithm based on branch-and-bound search, finds paths based on the mapped network generated from the atom mappings.

Despite the availability of several types of methods to understand and analyse metabolic networks, a simple and efficient method with minimum input for performing large-scale analyses is still lacking. Also, the current methods to identify pathways between the source and target molecules are restricted to smaller networks and do not enumerate long pathways, thereby restricting the scope of analyses. Although few of these algorithms handle cyclic and branched pathways, they require additional information such as atom transfer, which may not be readily available from the semi-curated metabolic networks. Further, we find that many of these tools are web-based, and do not lend easily to large-scale analyses; a large fraction of these tools are also currently inaccessible (Supplementary Table S1).

Thus, there is a need for a method to find pathways, which (a) requires only the topology of the reaction network (rather than stoichiometry and atom mapping), (b) is simple and scalable to metabolic networks (especially those comprising more than one organism), (c) efficiently handles cyclic and branched pathways, and (d) examines multiple alternate routes of conversion. To this end, we developed MetQuest, an efficient and scalable graph-theoretic algorithm, which exhaustively identifies all the pathways, or “sub-networks” between a given set of source and target metabolites using the input reaction network. Due to the nature of implementation, we are able to successfully identify branched and cyclic pathways. In comparison to the other algorithms, we observe better results, in terms of the completeness of the output pathways. Since we perform exhaustive enumeration, we not only recover the well-known pathways such as glycolysis and amino acid biosynthesis, but also identify several diverse pathways such as those involved in biodegradation of pollutants such as catechol. Further, because of the scalability of our algorithm to operate on larger metabolic networks, we are also able to demonstrate several metabolic exchanges happening in known natural and synthetic microbial communities.

## 2 Methods

In this section, we present a detailed overview of our algorithm, broadly divided into two phases. The first phase involves a *guided* breadth-first search (BFS) of the metabolic network, to identify all metabolites that can be reached from the source. In the second phase, we design a dynamic programming algorithm, which solves the non-trivial problem of assembling reactions into pathways to produce the metabolite of interest.

The primary aim here, is to find different sets of reactions that would be involved in the conversion of source(s) to target(s). Specifically, we find the pathways that comprise at most *β* reactions each, to enable this conversion. We define a pathway or sub-network as a set of reactions that are *together* necessary to convert a given set of source compounds to one or more target molecules. MetQuest seeks to efficiently traverse a metabolic network with branched and cyclic pathways and identify sub-networks for a given pair of source–target sets.

The input to MetQuest is a directed bipartite graph G derived from a given metabolic network, a specified set of source (*ϒ*) and a set of target (*T*) metabolites, a set of seed metabolites (*S*), a number *β* denoting the maximum number of reactions in any output sub-network. Any given metabolic network can be represented as a directed bipartite graph *G*(*M, R, E*), where *M* is the set of metabolites in the metabolic network, *R* is the set of reactions and *E* is the set of edges. Directed edges connect reactant metabolites (*m_i_* ∈ *M*) to a reaction node (*r_j_* ∈ *R*) or a reaction node to product metabolites. Such a representation disallows invalid conversions as may be interpreted from substrate graphs and helps in generating valid paths with biologically *meaningful* conversions [12].

### 2.1 Phase 1: Guided BFS

BFS is a classic graph traversal technique that visits all nodes of a given graph, starting at a source node, in a breadth-first fashion. BFS employs a queue of vertices, where newly discovered vertices are enqueued, to be processed at a later stage. A complete description of the BFS algorithm can be found elsewhere [8]. We modify the standard BFS by *guiding* it, based on the availability of precursor metabolites. Starting with the set of seed metabolites S, the algorithm first finds all the reactions from set *R*, whose precursor metabolites are in *S*. Such reactions are marked “visited” and added to the visited reaction set *R_v_*. The metabolites produced by these reactions, *m_c_*, are then added to *S*. The traversal continues in a *breadth-first* manner, incrementally adding triggerable reactions to the BFS queue. The expansion stops when there are no further reactions that can be visited. During the expansion, a reaction node is labelled as *stuck*, if it does not (yet) have the necessary precursors in *S*. Such reactions are automatically triggered if the precursor metabolites are produced at any later stage. A formal description of the algorithm can be found in Supplementary Algorithm S1. The traversed graph consists of all reactions that can be *visited*.

At the end of the traversal, we obtain the scope *M_s_* ⊆ *M* of *S* and the set of visited reaction nodes, *R_v_*. We also obtain the minimum number of steps to reach any metabolite m and reaction *r*, starting at the seed metabolite set *S*, denoted as *ℓ_m_* and *ℓ_r_*, respectively. We note that the value *ℓ_m_* may not indicate the exact minimum number of reactions required to produce a metabolite. Instead, we only leverage it for algorithm optimisation (see Supplementary Methods §4.1). The scope *M_s_* comprises all metabolites that can be produced from the seed set *S*, in the given metabolic network. This process resembles the ideas of network expansion [18] and forward propagation [1] reported earlier. The former method seeks to identify the synthesising capacity of metabolic network using the input seed set of metabolites. The latter aims to identify the minimal precursor sets of seed metabolites that are required to produce a given set of target metabolites.

This step in our algorithm also derives the information about the scope of metabolites from the given metabolic network; it however distinguishes itself by making a systematic note of the visited and stuck reaction nodes, which are exploited at a later stage for efficient and exhaustive enumeration of biosynthetic pathways.

### 2.2 Phase 2: Generation of Sub-Networks

MetQuest uses a recursive dynamic programming formulation for computation of sub-networks. However, it avoids repeated recursive calls by *memoising* the precomputed values. The algorithm steps are listed below and the pseudo code can be found in Algorithm 1. Our algorithm maintains a table of size *M_s_* × *β* and initialises all the entries to ⊥, indicating that we do not (yet) know about the sub-networks for every metabolite. Recall that *M_s_* and *β* denote the *scope* of seed metabolites and the maximum number of reactions we allow in a sub-network (“size cut-off”) respectively. We start filling the table entries by first considering the seed metabolite set *S*. For every seed metabolite *m* ∈ *S*, the entry in corresponding cell *Table*(*m*, 0) = ∅, indicating that no reaction is required to produce it; entry against other metabolites remain as ⊥ (Lines 1–5).

#### Algorithm 1 MetQuest

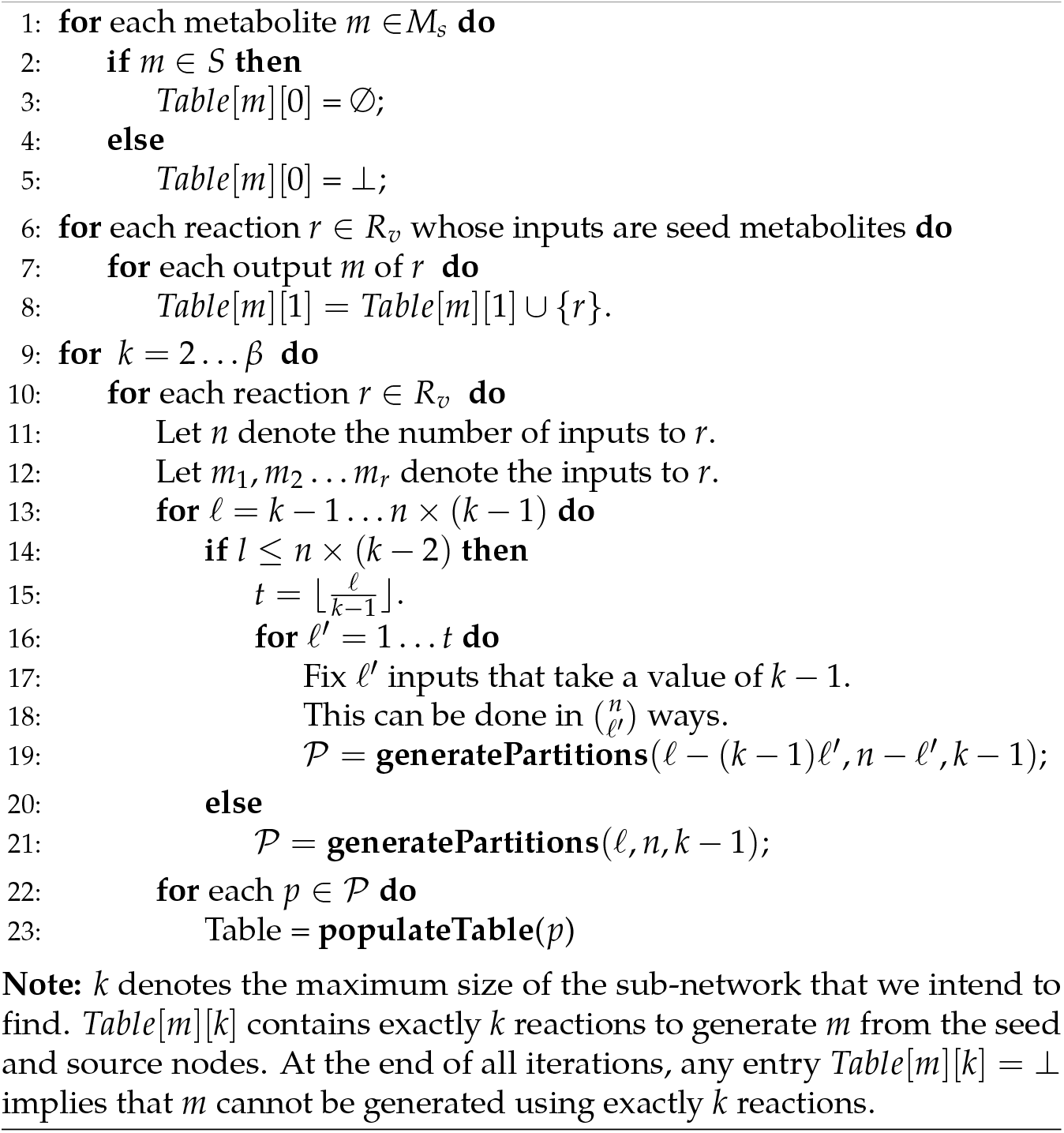

#### 2.2.1 Algorithm Steps

1. In the first iteration, we find all the reactions *r* ∈ *R_v_*, whose inputs are only seed metabolites. For every product metabolite *m* produced by *r*, we fill *Table*(*m*, 1) with the reaction that produced *m* (Lines 6–8)
2. We now propose that a pathway whose size *k* = 2,…, *β*, consisting of a triggerable reaction *r* ∈ *R_v_*, can be constructed by merging the pathways that generated the reactants of *r*. Let *n* denote the number of inputs to reaction *r* and *m*_1_, *m*_2_,…, *m_r_* denote the metabolites participating in *r*. In order to account for the branched pathways, it is necessary to generate all possible combinations of pathways producing these metabolites. However, evaluating all these combinations becomes computationally challenging since the metabolic networks are branched and there may be multiple combinations to be evaluated. To achieve this efficiently, we generate partitions using *n* non-negative integers, which sum from *k* − 1 to *n* × (*k* – 1) (see **generate Partitions**, as detailed in Supplementary Methods §4.2). By assessing the partitions for all the values till *n* × (*k* − 1), we guarantee the generation of all the pathways for the metabolites in the scope *M_s_* (Supplementary Methods §4.3). In order to achieve this sum, we choose from the set of integers whose value is between *ℓ_m_* and *k* − 1 (and not 0 and *k* − 1, as a part of optimization).
3. We generate such partition of numbers for all the reactions *r* ∈ *R_v_*, whose *ℓ_r_* ≤ *k*
4. Using these partitions of numbers, we fetch the sub-networks of a particular length (dictated by the numbers in that partition) for the input metabolites *m* from the corresponding cell entry in the table.
5. At a time, we consider one sub-network 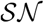 of every metabolite *m* in the input of *r*.
6. We then perform a union of these sub-networks for all the input metabolites *m*, and finally add the reaction *r* to it. We denote this resultant sub-network as 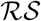. These operations are carried out by the function **populate Table** (Algorithm 2).
7. Against every output metabolite *m* of *r*, based on the value obtained in 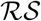, we add the entry to corresponding cell.
8. Since there can be several sub-networks producing these metabolites, we repeat the steps 5, 6 and 7 for all the entries.
9. The final reaction set will be a union of all the reaction sets that have been found for this metabolite.
10. We divide the process of generating partitions for numbers into two major stages, in order to avoid repetitions and optimise on running times (Lines 13–19 and Lines 20–21). More details can be found in Supplementary Methods §4.1. We note that the output pathways for every metabolite are generated by considering the sub-networks of *all* the reactants of a reaction. Due to this, any pathway generated by our algorithm is *complete*, i.e., it comprises of all the reactants required by the reactions constituting the pathway. Further, since for any metabolite, we perform the union of sets of reactions, we avoid repeated generation of the same reaction sets. Due to this, we automatically report only the first occurrence of cyclic pathways. The demonstration of our algorithm on a toy-network and a cyclic pathway can be found in Supplementary Methods §4.4 and §4.5 respectively. Further implementation details and the formal proof of correctness for our algorithm can be found in Supplementary Methods §4.6 and §4.7, respectively.

##### Algorithm 2 populateTable(p)

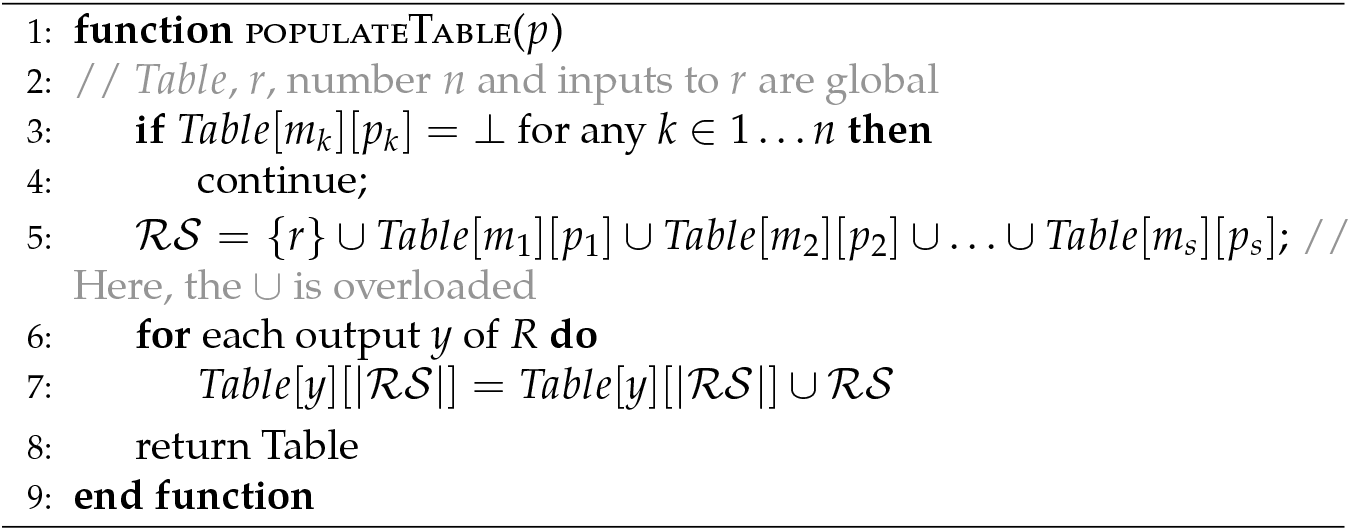

## 3 Results

In this section, we show that our algorithm can recover sub-networks of different types, including the well-known glycolysis and amino acid synthesis pathways. We also compare our output sub-networks with those generated by other algorithms, and demonstrate that MetQuest produces better results since it generates *complete* pathways. Further, we showcase the ability of MetQuest to identify diverse pathways involved in biodegradation of an important industrial pollutant. Finally, we show that MetQuest scales well to large networks, and is therefore a powerful tool to predict and understand metabolic exchanges in microbial communities.

### 3.1 MetQuest exhaustively identifies multiple pathways in metabolic networks

Genome-scale metabolic networks catalogue numerous metabolic pathways happening inside a cell. Identifying these pathways helps us to better understand the level of redundancy in the cells for producing key metabolites. To this end, we applied MetQuest on well-curated networks to identify and understand pathways between different substrates and products.

#### 3.1.1 Central carbon metabolism

We identify well-known biochemical pathways such as glycolysis using the genome-scale metabolic model of *E. coli i*JO1366 [38]. We constructed the corresponding directed bipartite graph, consisting of 12,974 edges and 5,659 nodes, and specified glucose as the source node and pyruvate as the target node. We included the co-factors, co-enzymes and the energy currencies in the seed metabolite set *S* (Supplementary Table S2) and found all the sub-networks within a size cut-off *β* of 15.

We successfully recovered the well-known glycolysis pathway and, in addition, due to the exhaustive enumeration, we could also identify 12,686 paths from the seed and the source set of metabolites to the target of varying lengths. Of all these paths, 6,175 of them use glucose as the source, while the rest of the paths comprise compounds formed by seed metabolites reacting amongst themselves.

These observations raise an important question: “Why should so many pathways potentially exist inside the cell in addition to glycolysis, which is the most preferred pathway?” To answer this question and understand the differences between each of these pathways, we studied the similarity between the obtained sub-networks, using the Jaccard index. Of ≈ 1.9 × 10^7^ combinations of sub-networks analysed we find only 2,858 (0.0149%) subnetworks are similar with Jaccard values between 0.93 and 0.98, while 8,992,712 (47.17%) sub-networks are much different with Jaccard values between 0.03 and 0.13 (Supplementary Figure S1).

The former set of sub-network pairs, with higher Jaccard values can be understood in light of the presence of multiple transporters, which are either used to transport compounds from the environment or between different compartments of the cells. These transporters function under varied environmental conditions, where they employ different mechanisms to transport glucose inside the cell [20]. The latter set, with lower Jaccard values, point towards the existence of alternate pathways that produce identical precursor metabolites to synthesise pyruvate. These predominantly include methyl glyoxal pathway, Entner-Doudoroff pathway and pentose-phosphate pathway. A few of these alternate pathways were also shown to outperform the canonical pathways under different sets of physiological conditions [9].

#### 3.1.2 Amino acid biosynthesis

We also determined sub-networks between different metabolites in a compartmentalised model of *S. cerevisiae* iMM904 [33]. As there are additional transport reactions between the compartments, we used a higher cut-off *β* = 30 and found the sub-networks between D-Glucose and amino acids such as L-phenylalanine. We assumed a seed metabolite set with 42 metabolites (Supplementary Table S3), consisting of the co-factors and co-enzymes in different compartments, and found sub-networks to 638 metabolites (|*M_s_*| was 681). Specifically, we identified 1,293 sub-networks from the source to target of varying lengths. From the sub-networks, we find that many of these amino acids are derived from the intermediates of the central carbon metabolism (Supplementary Results §5.1). Also, the sub-networks producing aromatic amino acids involve metabolites produced in different compartments of the cell, thereby requiring longer steps for conversion (Figure 1).

**Figure 1.**
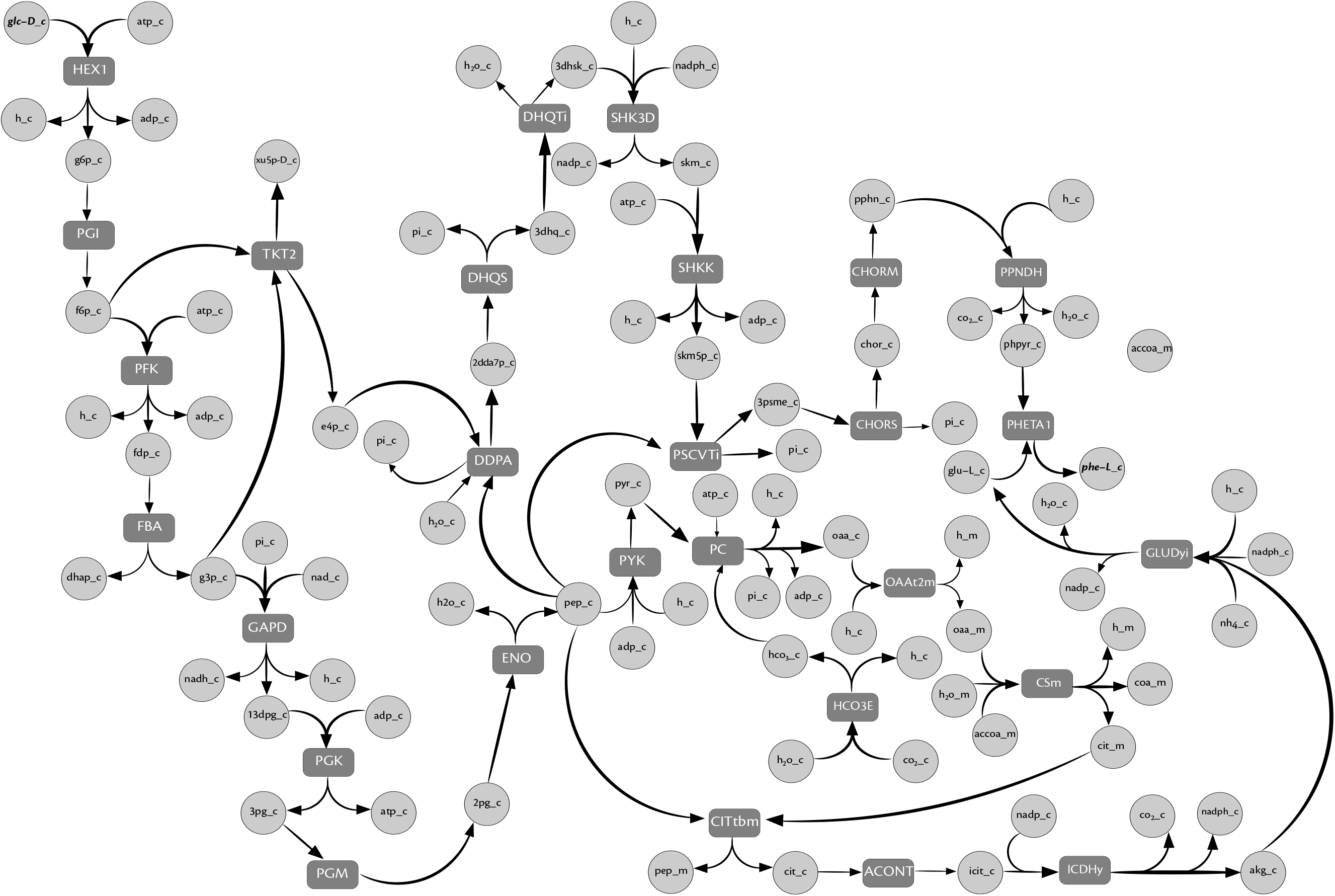
Sub-network with a size cut-off *β* of 28 between D-glucose and L-phenylalanine in *S. cerevisiae i*MM904. Dark gray rectangles represent reactions, light gray circles represent metabolites. The nomenclature of reaction and metabolite names are consistent with *i*MM904 genome-scale metabolic model. The source and the target metabolite is glc-D_c and phe-L_c respectively (shown in boldface and italics). We note that the synthesis of L-phenylalanine involves several metabolites from different compartments. The reactions in this pathway and the seed metabolites can be found in Supplementary Results §5.1.

### 3.2 MetQuest also uncovers metabolic dependencies

We also investigated if there were specific *centrally located* amino acids in the metabolic network, which play a crucial role in synthesising other amino acids. Towards this end, we constructed a bipartite graph using the genome-scale metabolic model of *E. coli i*JO1366 [38]. We initialized our algorithm with a set of seed metabolites consisting of essential cofactors and D-glucose as the source node (Supplementary Table S4), and generated all the subnetworks for a cut-off *β* = 20. We analysed the sub-networks for synthesis of different amino acids, to determine if there were considerable differences in their sizes. We studied this by progressively adding amino acids (one at a time) to the seed set, and determining if there were significant reductions in the size of sub-networks producing other amino acids. In each iteration, the amino acid that is produced via the fewest steps is added to the seed set. At the end of all iterations, we found amino acids such as L-Glutamate to be the key players in amino-acid biosynthesis. This is because these amino acids, when added to the seed metabolite set, considerably reduce the size of sub-networks producing amino acids such as L-serine and L-alanine (Supplementary Results §5.2). These results reiterate the central role of L-glutamate in the metabolic network as shown previously [3].

Further, we determined the sub-networks to these amino acids, and found one of our sub-networks producing L-histidine, matches the one reported in LumpGEM [2], which uses a Mixed-Integer Linear Programming based approach to identify reaction sub-networks. We could, however, not find any sub-network producing L-cysteine, L-methionine and L-tryptophan. This is because the former two amino acids need special co-substrates such as *S*-Adenosyl-L-methionine for their synthesis. These co-substrates, could neither be synthesised by this network from the given seed metabolites, nor were a part of the input seed provided. The latter amino acid, L-tryptophan is derived from the reaction of L-serine and other compounds such as indole, C’-(3-Indolyl)-glycerol 3-phosphate. These examples showcase the ability of MetQuest to exhaustively identify not only long pathways, but also uncover the metabolic dependencies in the metabolic network.

## 3.3 MetQuest reveals diverse paths for catechol biodegradation

Phenols and catechols are among the primary pollutants in the effluent from several industries, such as the chemical, textile and steel. So far, many studies have used *Pseudomonas putida* for bioremediation of such pollutants [24]. We sought to identify the pathways that render *P. putida* unique and tailored for this application. To this end, we investigated pathways involved in the metabolism of such compounds, which are otherwise harmful to many other micro-organisms. Further, we explored if the breakdown of catechols could directly lead to energy producing pathways. Hence, we identified all the pathways that start from catechols and lead to the intermediates of the tricarboxylic acid (TCA) cycle. Since catechols require complex mechanisms to be degraded to smaller intermediates, we used a higher cut-off *β* of 25. Using this, we found 825 different paths of lengths up to 25 that degrade catechols to fumarate. Interestingly, our algorithm could identify the two most different mechanisms of ortho- and meta-cleavage used by *Pseudomonas putida* to degrade catechols to the intermediates of the TCA cycle (Figure 2), as reported by [15]. These sub-networks can be found in Supplementary Results §5.3.

**Figure 2.**
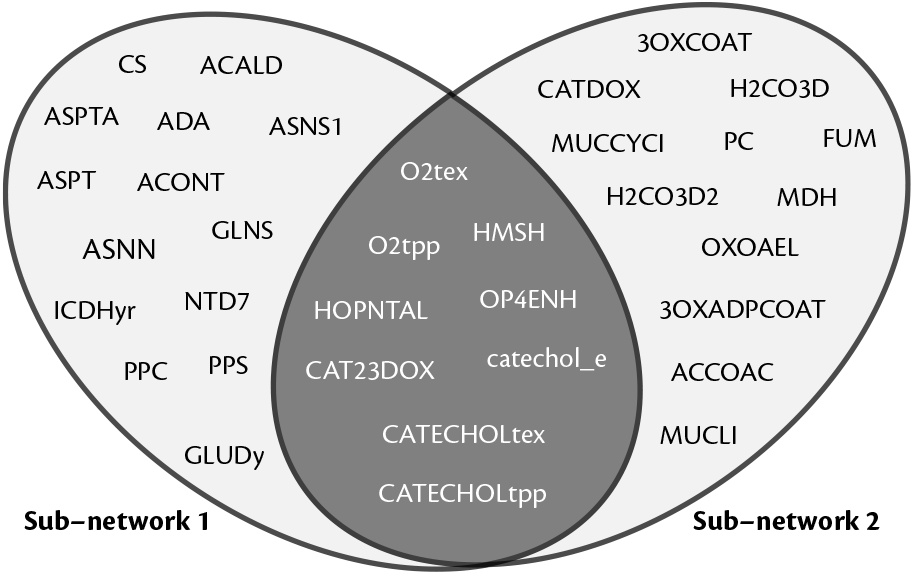
Overlap of the two most different pathways that degrade catechols in *Pseudomonas putida*. The two ovals represent reactions present in the two most different sub-networks identified. The intersection of the reactions found in the two sub-networks (shaded dark grey) pertain to the uptake of catechol. All the reaction identifiers follow the nomenclature from [36]. These reactions can be found in Supplementary Results §5.3.

## 3.4 MetQuest excels in comparison with other algorithms

To benchmark our algorithm, we compared our results with those obtained from some of the already existing path-finding methods such as FMM [7], ReTrace [40] and the pathways generated in ATLAS database [16]. Specifically, we determined paths/pathways between different source and target molecules, for multiple size cut-offs *β*. We constructed the bipartite graph corresponding to all reactions in the KEGG database [21]. For all the test-cases, we used the restricted compounds listed in [30] as the seed set of metabolites.

We find that MetQuest performs better, in terms of the number and the completeness of the pathways generated, i.e., the output sub-networks are *complete*, in that they have all the reactions necessary to produce every reactant in that pathway. From the sub-networks (Table 1), we observe that the smaller pathways of length 2 completely match with those generated by the other algorithms. However, in many cases, we identify longer pathways, since these involve metabolites generated by branched pathways. It is interesting to note that our algorithm was able to correctly identify the already reported pathway between C00418 (Mevalonic acid) and C16028 (Amorpha-4,11-diene) [29], which was not identified by the other algorithms. All the sub-networks reported in Table 1 can be found in Supplementary Results §5.4.

**Table 1.**
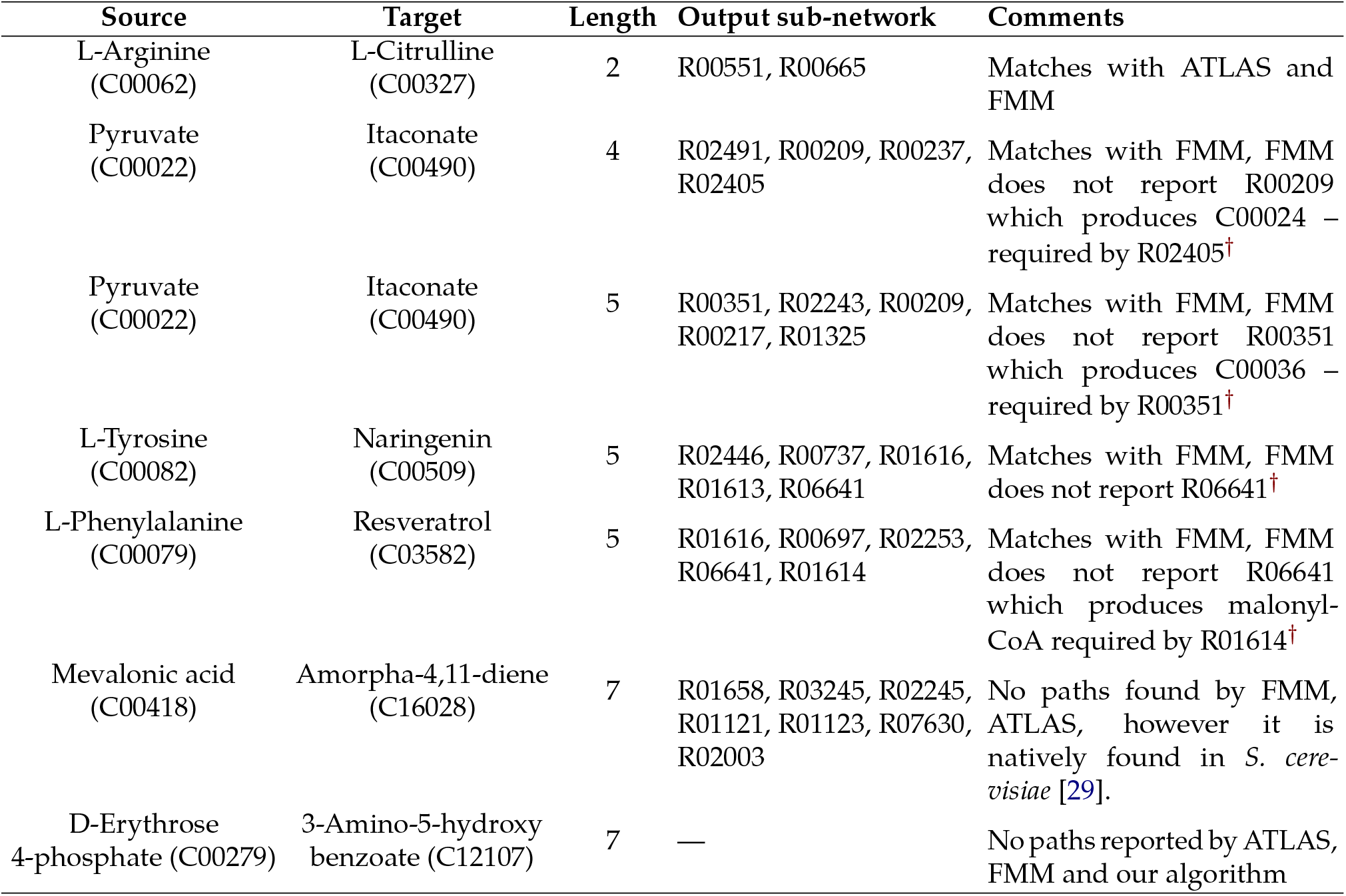
The sub-networks produced between different source and target molecules for a given length. These results were obtained on the graph constructed using reactions in the KEGG database. We denote reactions and compounds using their respective KEGG IDs. It is interesting to note that our algorithm is able to recover the well-known pathway reported in *S.cerevisiae*, while the other algorithms fail to do so. Also, there exists no pathways of length 7 between C00279 and C12107, which is being consistently reported by all the methods. More details about the sub-networks can be found in Supplementary Results §5.4.

We also tested the performance of MetQuest by finding sub-networks of varying size cut-offs (10–25) between glucose and other key metabolites involved in the central carbon metabolism on several genome-scale metabolic networks of different sizes (Table 2). We used a uniform set of seed metabolites consisting of essential co-enzymes and co-factors for all the simulations (Supplementary Table S5) and determined the sub-networks of sizes 10, 15, 20 and 25. During each simulation, we measured the time taken and the number of sub-networks generated for the target metabolites pertaining to the given size cut-off *β*. We carried out all the simulations on an Intel Core *i*7-2600 Desktop with 24GB RAM, running Ubuntu 14.04. Although we see an increase in the simulation time as the model size and the cut-off *β* increases (Supplementary Figure S2), MetQuest never fails to generate pathways, even at higher values of *β*.

**Table 2.** Number of sub-networks found for each target of given cut-off *β* from the seed metabolite set S (which also includes the source metabolite). Note that there are many sub-networks of length 20 for pyruvate in all the models but there are no sub-networks of length 10 that produce oxaloacetate in most of the models.

| Targets | Pyruvate |  |  |  | Oxaloacetate |  |  |  |
| --- | --- | --- | --- | --- | --- | --- | --- | --- |
| Models / Sub-network length | 10 | 15 | 20 | 25 | 10 | 15 | 20 | 25 |
| *Escherichia coli* core model | 14 | 142 | 2358 | 226 | 6 | 111 | 603 | 251 |
| *Lactococcus lactis* | 1 | 530 | 1673 | 1908 | No paths | 81 | No paths | 543 |
| *Pichia stipitis* iBB804 | No paths | 425 | 131 | 453 | No paths | 65 | 267 | 790 |
| *Saccharomyces cerevisiae* iMM904 | 3 | 1 | 1047 | 475 | No paths | 65 | 1423 | 2514 |

## 3.5 MetQuest scales well to large metabolic networks

Beyond individual networks, it is also interesting to identify sub-networks in communities of organisms and understand their metabolic interactions, which are known to play a critical role in determining the stability of microbial communities [14]. However, *community metabolic networks* are much larger in size, presenting many more challenges for path-finding. Analysing such large graphs of microbial communities demands a scalable and efficient algorithm, which requires only minimum information such as network topology. Further, the algorithm should be capable of identifying longer pathways to capture many metabolic exchanges happening within a community. The existing algorithms, such as OptCom [44] and cFBA [23], are based on constraint-based techniques, and require well-curated metabolic models. Also, they do not directly lend themselves to identify metabolic interactions in draft network reconstructions. To this end, we use MetQuest on *joint graphs* (metabolic networks) of multiple organisms and identify sub-networks between different metabolites. Specifically, we construct the *community bipartite graph* of a microbial community by considering a common extracelullar space through which metabolite exchanges can happen. Further, we analyse the sub-networks generated by MetQuest and show that it can correctly recover previously reported metabolic interactions.

### 3.5.1 MetQuest correctly predicts metabolic exchanges in a synthetic *E. coli* community

To study the metabolic interactions in *E. coli*, [43] performed computational analyses on genetically modified *E. coli* strains. Specifically, two different *E. coli* strains with knockouts of b2276 and b3708 genes were modelled together and the metabolic interactions were predicted. We simulated these gene knockouts by removing the reactions catalyzed by these genes, after considering the Gene-Protein-Reaction relationships. We then constructed the community bipartite graph and applied MetQuest to predict the metabolic exchanges between these two strains. We assumed *seed metabolites*, with glycolate as source (as given in [43]) and applied MetQuest to find all sets of sub-networks to every metabolite within the scope of glycolate, pyruvate and both these sources independently, with a size cut-off *β =* 20.

From the sub-networks generated, we determined the metabolites that can be potentially exchanged between the strains. We were able to recover the reported acetate and formate exchanges happening between the two strains. In addition, we observe that acetate (from Δ*b*2276) participates in the production of several important target metabolites in Δ*b*3708 such as L-glutamine, inosine and several other co-enzymes of central carbon metabolism. In addition, we also found other metabolites such as alpha-ketoglutarate, ethanol, acetaldehyde exchanged between the organisms, which can be potential candidates for experimental verification (Supplementary Results §5.5).

### 3.5.2 Predicting novel interactions in community metabolic networks

To illustrate the performance of MetQuest on a larger network, we computed multiple sub-networks in an experimentally demonstrated three-member microbial consortium [31], and determined the metabolic interactions. Towards this, we used the genome-scale metabolic models of *Clostridium cellulolyticum* (cc), *Desulfovibrio vulgaris* (dv), and *Geobacter sulfurreducens* (gs) from the Path2Models database [6]. We constructed the three-member community bipartite graph by connecting the genome-scale metabolic models through their common exchange reactions. We defined the seed metabolite set *S*, consisting of essential salts, co-factors, co-enzymes and a set of *tRNA* molecules. We specified source as cellobiose (as reported in [31]) and chose to determine sub-networks with a size cut-off *β* = 20, to all metabolites within the scope *M_s_* of seed metabolites. To determine the metabolic exchanges, we analysed every sub-network of 1,526 metabolites (*M_s_*) with a special focus on sub-networks involving exchange metabolites.

From this analysis, we found many sub-networks having at least one exchange reaction between the organisms. We observed that the acetate and ethanol exchanges proposed in [31] predominantly led to the production of amino acids such as L-serine, L-leucine, L-aspartate and L-valine (in gs) and L-threonine and L-glycine (in dv) respectively, which contribute to the biomass, thereby enabling a stable microbial consortium.

## 4 Discussion

With the increasing number of draft genome-scale metabolic reconstructions, there is a pressing need for efficient algorithms to analyse these metabolic networks and generate useful predictions. It is particularly important that these algorithms perform efficiently with minimum information such as the reaction topology, and also identify pathways in large (multi-)genome-scale metabolic networks.

To this end, we propose MetQuest, a scalable and efficient graph-based algorithm for identifying all possible metabolic pathways in genome-scale metabolic models. Specifically, we focus on exhaustively determining all alternate pathways (of a particular size) between a set of source and target metabolites. We also demonstrate the application of our algorithm by identifying multiple pathways involved in several important metabolisms such as the central-carbon metabolism and the amino acid metabolism. Further, we show its usefulness in determining several diverse pathways, which take part in catechol degradation. Besides its applications on the metabolic networks of individual organisms, MetQuest, due to its scalability, can be applied to study larger and more complex networks.

A number of graph-based methods to find pathways in metabolic networks have been developed so far. These methods convert the metabolic networks either into substrate graphs, bipartite graphs or hypergraphs. However, these methods are well-suited only for smaller networks and lower cut-offs. For instance, MetaPath [17] tries to back trace the pathway from the target to the source based on the information obtained from the scope calculation. Since the metabolic networks contain many branched pathways, identifying the routes of conversion by back-tracing becomes computationally expensive, and can break down rapidly at higher cut-offs. Another method, Rahnuma [32], tries to solve the problem of path prediction on genome-scale metabolic networks by abstracting these networks into hypergraphs. Rahnuma, based on depth-first search on hypergraphs, seeks to obtain pathways between two metabolites. However, the performance of Rahnuma on metabolic networks was demonstrated only for shorter path-lengths of 6, and it may not be applicable for identifying longer pathways, such as those involved in the biosynthesis of aromatic amino acids from simple sugars. Moreover, the pathways are computed based on a condition that the metabolite can be used only once in any pathway, which inherently fails to capture cyclic pathways. MetQuest, on the other hand, can identify branched and cyclic pathways, and can perform well on larger metabolic networks for much longer cut-offs, thereby extending its application to analyse microbial communities.

We have also demonstrated the major strengths of MetQuest: in multiple examples we considered, of individual metabolic networks and real-world microbial communities, MetQuest was able to identify multiple pathways and correctly predict the metabolic exchanges/interactions taking place.

MetQuest was able to correctly predict the acetate exchange happening between the two genetically modified *E. coli* strains. Further, MetQuest also predicted novel interactions involving amino acids such as L-cysteine in the microbial community reported in [31].

However, as with any modelling technique, MetQuest also has its limitations. Firstly, the predictions from MetQuest rely heavily on the quality of the underlying metabolic network. Specifically, few of the metabolic interactions predicted by MetQuest could be due to model artefacts. There may be other metabolic interactions happening between microbes, which could be absent in the metabolic network (gaps in the metabolic network). Due to this reason, such interactions may not be predicted by MetQuest. Further, our algorithm does not assign any weights to metabolites/paths. Although there are other methods like cFBA and OptCom, which make more quantitative predictions, they also demand better curated metabolic networks. We primarily see the use of MetQuest as a first-line investigatory tool, to cull down from a very large set of possible communities to a more tractable set, which may be studied further using constraint-based techniques or through wet lab experiments.

In sum, we report a novel and efficient algorithm MetQuest, which (i) rapidly enumerates all possible biosynthetic pathways from a given metabolic network, (ii) efficiently handles branched and cyclic pathways, and most importantly, (iii) scales well to very large networks such as those representing microbial communities. Further, our algorithm requires only the genome-scale metabolic networks, a set of seed and target metabolites and a size cut-off, as against thermodynamic or atom-mapping information, thereby rendering it a very potent tool to perform large-scale analyses. Overall, we believe that MetQuest is a very useful tool for exploring and understanding metabolic pathways in large-scale networks, to generate testable hypotheses for further experiments.

## Funding

This work was supported by the Indian Institute of Technology Madras grant ERP/1314/004/RESF/KARH to KR and the INSPIRE fellowship, Department of Science and Technology, Government of India to AR.

